# Movement Recognition via Channel-Activation-Wise sEMG Attention

**DOI:** 10.1101/2023.06.03.543591

**Authors:** Jiaxuan Zhang, Yuki Matsuda, Manato Fujimoto, Hirohiko Suwa, Keiichi Yasumoto

**Author notes:** Corresponding author (J. Zhang).

## Abstract

**Context:** Surface electromyography (sEMG) signals contain rich information recorded from muscle movements and therefore reflect the user’s intention. sEMG has seen dominant applications in rehabilitation, clinical diagnosis as well as human engineering, etc. However, current feature extraction methods for sEMG signals have been seriously limited by its stochasticity, transiency, non-stationarity. *Objective:* Our objective is to combat the difficulties induced by the aforementioned downsides of sEMG and thereby extract representative features for various downstream movement recognition.

**Method:** We propose a novel 3-axis view of sEMG features composed of temporal, spatial, and channel-wise summary. We leverage the state-of-the-art architecture Transformer to enforce efficient parallel search and to get rid of limitations imposed by previous work in gesture classification.

**Results:** We compared the proposed method against existing methods on two Ninapro datasets consisting of data from both healthy people and amputees. Experimental results show the proposed method attains the state-of-the-art (SOTA) accuracy on both datasets. We further show that the proposed method enjoys strong generalization ability: a new SOTA is achieved by pretraining the model on a different dataset followed by fine-tuning it on the target dataset.

## 1. Introduction

Studying neuromuscular processes in electrical stimulation is an important step toward understanding the myoelectric control mechanism in humans [1]. Surface electromyography (sEMG) is a diagnostic tool recording the activation of the muscle cells with millisecond resolution that is widely used for measuring and evaluating the electrical response associated with skeletal muscle activities [2]. Dated back to the 1960s, sEMG has been used as the capable signal to control simple prosthetic grippers with a single degree of freedom (DOF). In recent years, sEMG has also seen applications in computer games [3], virtual reality [4], human-machine interfaces (HMI) [5], robotic control [6]. But perhaps more importantly, it has been identified as a powerful tool in physiology such as assistant diagnosis in identifying abnormal gaits, Parkinson’s disease (PD) [7], and Cerebral palsy (CP) in children [8].

Successful sEMG applications require high-quality and informative signal recordings. Unfortunately, there are several non-trivial obstacles in the way of obtaining clear and informative sEMG patterns. For example, sEMG signals are transient, stochastic, non-stationary, nonlinear and non-predictable [9]. Noise contamination composes another problem and is usually unavoidable due to the nature of non-invasive signal acquisition from the surface of the skin. Depending on sEMG collecting equipment and protocol, different types of noise can be found within the signals such as motion artifacts or power line interference. The level of contamination of sEMG signals is dynamic and varies in realworld environments, introducing confounding variability to the signals and may uncorrelated the signals to the motions of interest [10]. Besides noises, muscles themselves are potential sources of confusion: a muscle is composed of many motor units (MU), and the *discharge* or *firing* process of each MU generates a *motor unit action potential* which is the sum of the contributions from individual fibers that compose the MU [11]. As a result, sEMG recordings always contain massive and redundant information, reliably extracting informative representations from such massive noise-involved and redundant signals is non-trivial and has been a pending challenge to physiological research as well as to any downstream tasks.

Conventional feature extraction on sEMG recordings emphasizes expert knowledge by manually selecting features, often the output of some statistical methods from spatial or spatio-temporal spaces such as spectrum, wavelet, etc [12]. If the hand-crafted feature spaces are well-established, superior performance can be expected. Indeed, classic machine learning methods such as Random Forest (RF) [13], k-Nearest Neighbor (kNN) [14], and support vector machine (SVM) [9] have recently been dominant in the community as they have shown promising performance on top of expert-made feature spaces. However, increasing evidence shows that these conventional methods can fail miserably for poorly or incompletely captured features in complicated tasks such as stochastic and transient bursts. This is because these methods represent only sensible decision-making given informative features, yet leaving how to build an informative feature space given complicated signal sources a pending question. In light of the aforementioned downsides of manual feature engineering, numerous studies have been devoted to building end-to-end pipelines; that is, methods that integrate feature extraction and decision-making into a single machine learning model for various downstream tasks.

The recently successful deep learning has provided a possible solution to the aforementioned problems and building superior end-to-end platforms [15]. Promising preliminary results have been shown on surface muscle signal processing problems. Theoretically, it is well-known that deep learning models with nonlinearity can approximate complicated nonlinear functions arbitrarily well, which stands as a sheer contrast to the conventional methods. Practically, variants of deep learning models have been developed for different use cases. For example, some works leveraged convolutional neural networks to capture local patterns and variations from sEMG as informative features [16]. However, though convolution can express channel connectivity of different muscle signals, the local inductive bias of CNNs inevitably leads to the loss of global information [17]. Another branch of work focused on modeling the time-series property of sEMG signals [18]. Recurrent neural networks (RNNs), such as Long Short Term Memory (LSTM), bi-LSTM, and Gated recurrent unit (GRU) can effectively summarize global information that varies with time [19]. However, the issue of variability in sEMG signals, caused by the transient and stochastic nature of the neural drive to muscles persists and can greatly perturb the performance of RNNs. While remarkable performance has been shown in language processing in which sentences have global structures, RNNs often struggle with temporal sEMG problems where local information such as the time-dependent spikes in sEMG are as important as global contexts such as the number of bursts or overall amplitude.

The nuances as the result of noise-contaminated muscle motor units apparently demand more sophisticated models capable of providing extra modeling power over existing architectures such as CNNs or RNNs: it is desired that both local variations and global summary of the conventional spatio-temporal domains could be extracted and displayed onto a new *feature domain* to serve as a better representation [20]. The state-of-the-art (SOTA) model on a variety of complicated tasks such as vision Transformer, [21, 22] comes to rescue by searching among all sEMG channels for those responsible for a specific muscle move [23]. The resultant novel time-frequency-channel representation offers a new paradigm for sEMG feature extraction: since human movements typically activate only a few number of muscles, the majority of sEMG channels are not responsible for that move. Taking as input the whole channel information as CNNs or RNNs can corrupt information since noise accumulates across channels as the model proceed sequentially. By contrast, the channel-wise parallel processing of the Transformer is capable of quickly finding the channels that are responsible for a specific move, hence providing more informative and less contaminated features.

Specifically, we propose a novel, better feature extraction method that combines conventional feature processing with a state-of-the-art deep learning model. For each channel, we first apply wavelet transform to extract distinct frequency features. Similar to its application in word embedding, the Transformer is leveraged to embed local spatiotemporal wavelets in a channel-wise manner and to globally search for specific channels responsible for a given muscle movement. In this work, we consider sEMG classification but the time-frequency-channel representation is generally applicable and we believe it is the key contribution of the paper. To verify the proposed method, we compare it against several well-established feature engineering methods and perform feature ranking to find what constitutes important features. We summarize the main contributions of this paper as the following:

- This work is the first approach to propose a novel 3axis feature representation for sEMG signals realized by a new architecture built upon the state-of-the-art Transformer model.
- We compare against conventional feature engineering methods and perform feature ranking to demonstrate the proposed pipeline enjoys strong interpretability besides superior performance.
- We successfully achieve new state-of-the-art classification results by the novel framework in different databases.

The paper is organized as follows: Section II provides a brief overview of relevant literature. In Section III, we present the dataset and feature extraction. The proposed architecture is developed in Section IV. Experimental results and different evaluation scenarios are presented in Section V. Finally, Section VI concludes the paper.

## 2. Related Work

In this section, we position our work in the literature by providing a brief review of related work in chronicle order to illustrate how sEMG models have evolved from the conventional statistical learning-based methods to recent architectures focusing on sophisticated deep models.

### 2.1. Statistical Learning Methods

It is vital to recognition as well as to any downstream tasks how the sEMG features are extracted and represented.

Conventionally, such representation extraction is done by manually selecting from the spatial domain and/or the temporal domain [24]. Different statistical methods are used for extraction, such as spectrogram transformation [25], entropy-based analysis [26], spike simulation [2], wavelet transformation [1], empirical mode decomposition [27], etc. Representative and informative features are the guarantee for the performance of subsequent learning. In the community, the current dominant trend is to have the classic statistical learning algorithms integrate the features and distill recognizable sEMG features for downstream tasks. For example, Tommasi et al. [28] proposed to utilize SVM to classify up to seven different hand movements by adaptation based on multiple pre-trained models. In [29], a classification task composed of 15 hand movements from persons with intact limbs and 12 from amputees was performed leveraging the Linear Discriminant Analysis (LDA) and SVM based on the autoregressive features. In [30], energy-based features were used with the nonparametric kNN for the classification of gestures using the sEMG recordings. In all these statistical learning methods, a key limitation is that the feature spaces could never be more expressive than the optimal spanning of the manually selected features. Manual design of relevant features is often incomplete, inaccurate, and insufficient for representing highly complicated patterns [4], say the nonlinear non-stationary variations.

### 2.2. Deep Learning Based Methods

Benefiting from deep architectures composed of nonlinear activation units, deep learning models have been shown to be capable of approximating complicated functions arbitrarily well. The community has seen an increasing number of research leveraging deep learning as an automated feature extractor to decode sEMG signals [31]. Moreover, deep learning enjoys additional benefits such as the ease of integrating decision-making module with the feature extractor into one end-to-end model. Such models can significantly benefit from the recent advance in computing power by deploying the model in parallel such as on the powerful graphics processing unit. Quivira et al. [32] introduced CNNs to the diagnosis of neuromuscular disorders to extract wavelet features for detecting and classifying anomalous sEMG signals. Their experiment showed promising results that real-time detection of neuromuscular disorders is possible. Wei et al. [33] proposed to utilize multi-stream CNN that takes as input and process segment of images in parallel to provide patch-wise analysis of images. Their result demonstrated state-of-the-art accuracy at the cost of 10-20% higher computational complexity than other architectures. More recently, [34] built an accurate regression model to predict hand joint kinematics from sEMG features by using RNN with LSTM cells to account for the non-linear relationship between sEMG and hand poses. They experimented with 7 users to test the performance of the proposal and the results showed the model was capable of accurately capturing simple hand gestures such as finger flexion. It is worth noting that, the mainstream SOTA deep learning models such as variants of CNNs or RNNs have a salient shortcoming for sEMG signals: the sequential features of sEMG signals tend to be ignored. This issue is especially noticeable in the much more time-consuming and unstable RNN-based models.

### 2.3. The SOTA Transformer

As one of the most important machine learning tasks, vision and natural language processing have witnessed a breakthrough [35, 36] due to the advent of Transformer [21]. Different from previous deep learning models, it features the novel self-attention architecture that allows for efficient parallel search in the local as well as global context. Specifically, the attention mechanism attends more to contextually salient factors without considering the information transfer of time step [37]. The novel architecture proves to be superior when the modeled objects exhibit local-global structures such as sentences or images. In physiology and health care, it has very recently been utilized to provide clinical support [38].

In 2022, Transformer has seen extensions to sEMG recognition [31, 39]. Specifically, in work [31] a novel convolutional vision transformer (CviT) with stacking ensemble learning was proposed for the fusion of sequential and spatial features of sEMG signals with parallel training. Their experimental results demonstrated the proposed CviT obtained better performance than most current approaches. In addition, the successful application of Transformer in sEMG-based movements classification provides a significant reference for the application of Transformer in other biological signals [31]. In [39], the transformer architecture was leveraged for decoding object motions in dexterous in-hand manipulation tasks using raw EMG signals input. Their new architecture Temporal Multi-Channel Transformers and Vision Transformers were shown to outperform RF-based models and CNNs in terms of accuracy and speed of decoding the motion.

It is worth noting that, these work focused only on leveraging the transformer on the frequency and/or spatial-temporal features, neither considered expanding the feature space via another view as our proposed method do. As will be shown in the experimental section, our proposed new representation brings a significant improvement over their methods.

## 3. Methods

In this section, we describe the pipeline and the details of the proposed method. We first explain the signal preprocessing in Section 3.1. Then, Section 3.2 and 3.3 introduce the Spatio-temporal transformation and its feature extraction methods in different channels. Finally, the framework construction of the proposed channel-wise Transformer is described in Section 3.4.

### 3.1. Pre-processing

Our preprocessing pipeline consists of as the first step a signal filter. Since sEMG recordings are typically contaminated with various types of noises from sensing, device, etc, we used the 3-order Butterworth bandpass filter within a 5-500 Hz setting for each subject signal. The second step is sample generation by a 200ms moving window with half-overlapping for sample segmentation following the work [40].

### 3.2. DTCWT Transformation

As previously mentioned, this work aims to recognize informative sEMG patterns by three-axis features (temporal-, spatial-, and channel-domain), hence, we first generally exploit sEMG temporal-spatial features. Wavelet transformation is one of the most well-known signal processing methods that can provide temporal-spatial features [41]. The determination of wavelet type, mother wavelet, and implementation techniques are the key components of the wavelet transform based temporal-spatial representations. While many wavelets and variations have been proposed, it is commonly perceived that dual-tree complex wavelet transform (DTCWT) is currently the best wavelet transform used for sEMG recognition [42]. Because DTCWT approximates the shift-invariance property and improves upon the aliasing effects. In this work, we adopt the DTCWT to first extract temporal-spatial features followed by applying coefficients to effectively represent global and local features from sEMG samples.

In the first step of the wavelet transform, the input signal, *x*(*t*) passes through two filters: high- and low-frequency. Their outputs are called detail and approximation coefficients, respectively, here, *Ψ*_*h*_(*t*) and *Ψ*_*g*_(*t*) are real-valued wavelets and the complex-valued wavelet *Ψ c* (*t*) is obtained as

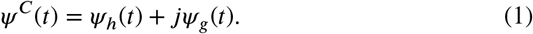

where the right items need to first calculate by the real component of wavelet coefficient *dl*_(*Re*)_(*k*) and approximation coefficient *a*_*j*(*Re*)_(*k*):

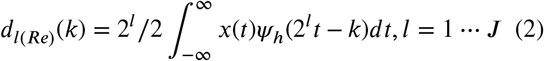

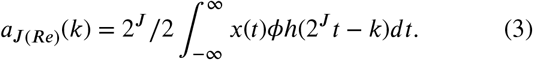

where *j* is the index of the scale factor *J* which controls the frequency content. Therefore, the output of DTCWT can be expressed by making a summation of the output trees.

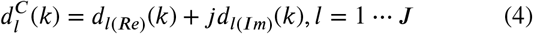

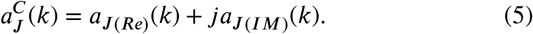

Consequently, the 1-D signal parts are obtained by replacing *Ψ*_*h*_(*t*) and *ϕ*_*h*_(*t*) with *Ψ*_*g*_(*t*) and *ϕ*_*g*_(*t*), respectively.

### 3.3. Spatio-Temporal Feature Extraction

Following the preprocessing of sEMG signals, the sEMG time series is changed to the time-frequency domain. This step is followed by feature extraction: a spatio-temporal vector (*X*_*ST*_) is generated by the following 18 feature extraction methods for each channel in each sample. The details of 18 feature extraction methods can be found in Table 1. After feature extraction, we concatenate all channel-wise vectors to construct a feature matrix for each sample denoted as *X*_*ST*_ *∈ ℝd*×*c*, where *D* represents the number of extracted spatial-temporal features and *C* the number of channels. We consider these features have some functional overlap, such as, the ‘Standard deviation’ and ‘Difference absolute standard deviation value’, and the redundant features may lead to noise to the sEMG recognition as well as decrease the performance. Hence, a deep feature ranking method is introduced to this work to refine the feature vector construction. The details will be shown in Section 3.4.2.

**Table 1.** 18 features extracted from DTCWT used in this paper and their brief explanations.

| ID | Method | Explanation |
| --- | --- | --- |
| *F1* | Mean Absolute Value | the mean absolute value of signals |
| *F2* | Standard Deviation | standard deviation of signals |
| *F3* | Skewness | 3rd moment of signal distribution |
| *F4* | Kurtosis | 4th moment of signal distribution |
| *F5* | Zero Crossing | #the instantaneous point at which there is no voltage present |
| *F6* | Average Energy | the squared sum of the sample values in the time window |
| *F7* | Waveform Length | the cumulative length of the waveform over the sample segment |
| *F8* | Maximum Fractal Length | the maximum length of fractal dimension feature |
| *F9* | Integrated EMG | the area under the curve of the rectified signal |
| *F10* | Root Mean Square | the square root of the arithmetic mean of the square values |
| *F11* | Log Detector | the estimated value of contractility |
| *F12* | Log Difference Absolute Mean Value | log difference of two independent values |
| *F13* | Coefficient of Variation | a standardized measure of the dispersion of frequency distribution |
| *F14* | Difference Absolute Mean Value | difference of two independent values drawn from a distribution |
| *F15* | Difference Absolute Standard Deviation Value | 1st difference of the dispersion in a set of data |
| *F16* | Difference Variance Value | difference variance value of wavelength |
| *F17* | Enhanced Mean Absolute Value | the estimated value of the exerted force |
| *F18* | Simple Square Integral | the quadratically integrable function |

### 3.4. Channel-wise Transformer

The extracted Feature matrix is fed into a deep-learning model for downstream tasks such as classification. However, the feature output should be conventionally compressed, e.g., reshaped to a vector for matching the input size of the fully connected layer. Such compression would destroy the independence between time-frequency and channel domains. Hence, a novel deep learning method, Transformer, that individually processes and summarizes the dependency is called for [22]. The transformer creates a set of sub-networks to handle different feature sub-spaces (rows or columns in the feature matrix). Relevant features among these sub-networks are then found by the well-known attention mechanism [43]. Introducing the Transformer to sEMG recognition naturally suits to our proposal for processing and tracking individual contributions from the sEMG channels. Each channel index vector *C* of *X*_*ST*_ is taken as input to a sub-network for retaining the feature space.

#### 3.4.1. Channel Sequence Embedding

In detail, for each input *X*_*ST*_, we first embed each column to a higher dimensional space to keep channel independence. We split the channels for independent linear projection. Each linear projection operation functions to spatiotemporal feature space that projects the original feature space with *d* dimension to a richer space of dimension *d*′. A channelwise sequence (*S*_*ch*_) is subsequently generated as the input to Transformer. Inspired by [22], an extra class token *S*_*Cls*_ is inserted at the beginning of each sequence *S*_*ch*_. As data pass through the model layers, *S*_*Cls*_ becomes a new feature summarizing the global relevance of channels for downstream decision-making. The above procedure can be described mathematically as:

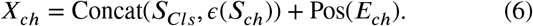

where *∈*(*·*) is the above mentioned channel-wise linear projection. Since the Transformer abandons the sequential order (positional index) of the input, a sequence of positional encoding with the same size of *S*_*Cls*_ is generated and used for learning the positional importance of each element of *s* during the model training. As a consequence, the sequence *X*_*ch*_ is the final input fed to the Transformer model servers to make the downstream decision.

#### 3.4.2. Feature Space Construction via Attention Mechanism

As shown in Figure 3, the Transformer consists of two layers: an *attention layer* and the *multi-layer perceptron* (MLP). The attention layer calculates the pair-wise relevance of different patches (sEMG channels) in *X*_*ch*_ and maps the relevance to the ground truth with three matrices: query *Q*, key *K*, and value *V*. These three matrices are the projected feature space of *X*_*ch*_, and the attention/relevance based on these matrices is computed as:

**Figure 1:**
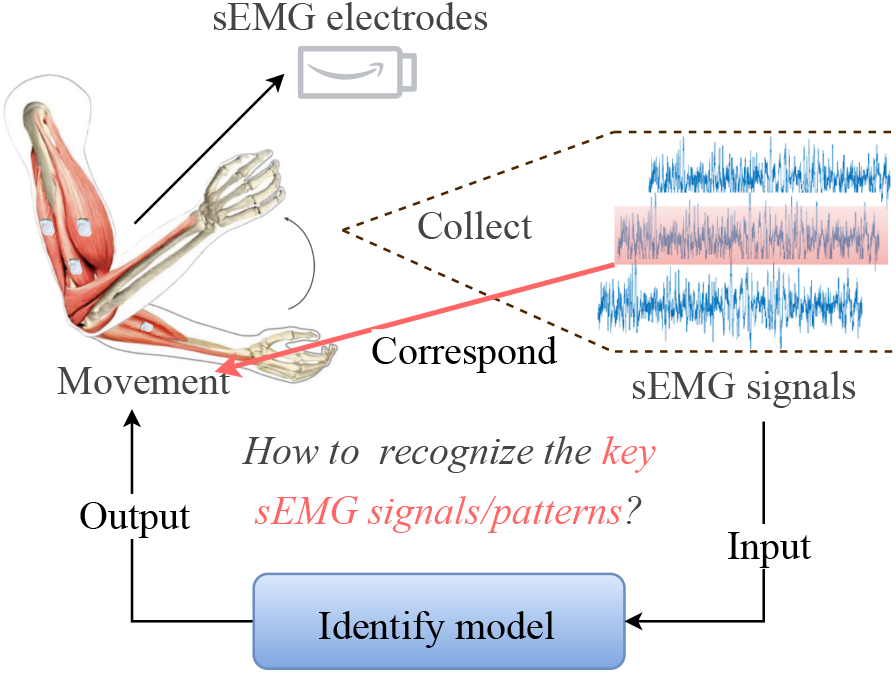
The key question in this work: *what is the right information domain for movement recognition by sEMG?* We would give an answer here: *the muscle-activation-wise sEMG channels*.

**Figure 2:**
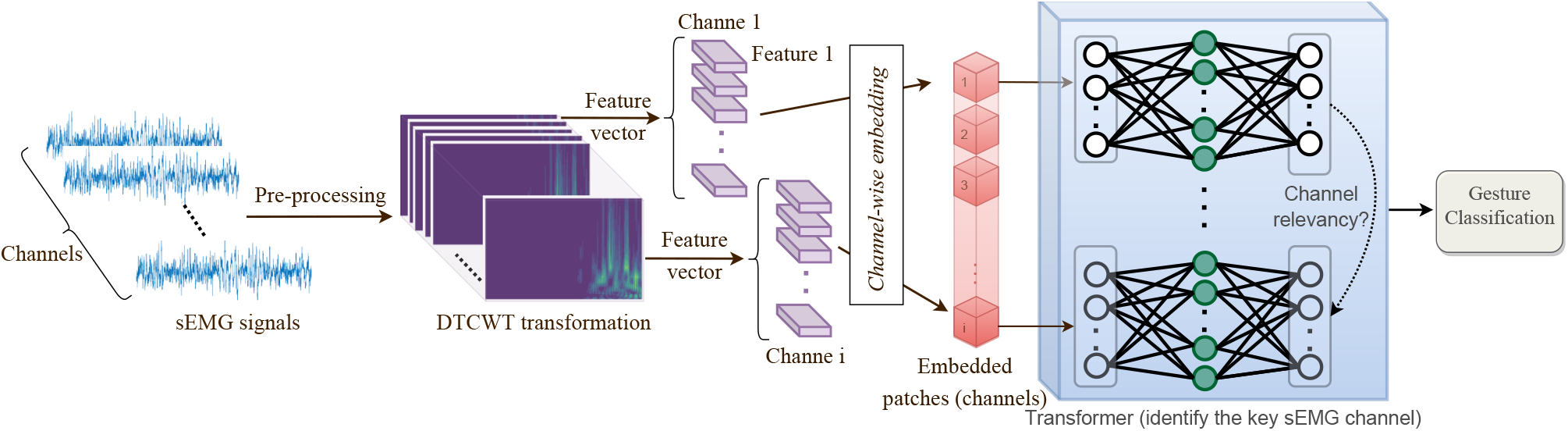
Overview of the proposal. The experiment of this work was done with multi-channel sEMGs. Each 200ms sEMGs episode is firstly converted to a time-frequency wavelet transformation. Then, different feature extraction methods are applied to this transformation and a set of feature vectors corresponding to different channels is generated. These feature vectors are embedded in a patch sequence. Each patch represents one feature vector for each sEMG channel is projected to a high dimension by using a patch-wise fully connected layer. Afterward, the augmented patch sequence is fed to the channel-wise Transformer model which can be viewed as containing a set of sub-networks. After training, a relevance calculation is conducted by the model to find the channel relevance for different sEMG recognition targets, e.g., different gestures.

**Figure 3:**
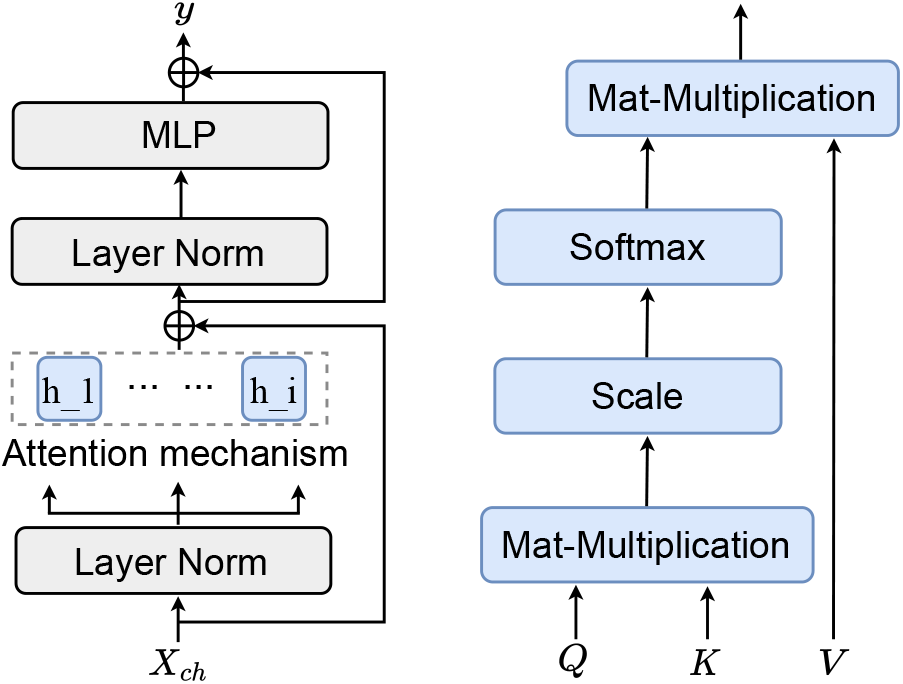
Attention mechanism and the compute flow of Transformer.

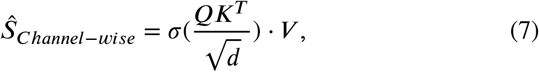

where (*·*) denotes a patch-wise softmax function. 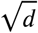 is a normalization operation that is applied to each *Q*-*K* computation to control the max-min range of the attention value. The resultant matrix ?^ records the attention score in different sEMG channels calculated by weighting the relevance resulting in *σ*(*·*) to each row of *V*.

Afterward, the normalized output of the multi-head attention layer is fed to the MLP layer. The activation function for the MLP layer is the Gaussian error linear units (GELUs) referring to [22]. When *s* iteratively passes through all layers of the Transformer during the training phase, *S*_*Cls*_ has summarized the channel’s relevancy information for different classes. This token is separated from the sequence of output and fed into a softmax layer to make the final classification decision. To achieve the feature ranking, we introduce variational dropout regularization [44] to the MLP layer. Because the MLP is the feature-wise MLP that functioned to different spatio-temporal features in each channel. The importance calculation and ranking are conducted by a max-min game of the model training: minimizing the prediction error of classification while maximizing the dropout rate for the rest of the unimportant features. Mathematically, given features ?, a variational mask distribution 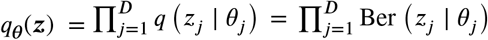 is estimated a fully factorized distribution [44]. Hence, the feature-wise dropout rate *θ*_*j*_ corresponds to the importance of the *j*-th feature of spatio-temporal features.

#### 3.4.3. Decision Making Module of Transformer

In practice, the *attention mechanism* is implemented in parallel. Specifically, similar to the multi-channel of CNNs, attention mechanism can be extended to a set of projected attention implementations (the well-known *multi-head* of Transformer). The multi-head attention prevents losing the manifold expression of the features. At the beginning of each building block, *h* (the number of the heads) sets of *Q* and *K* are generated and mapped by the linear projection. Then, the attention implements *h* times in parallel to calculate relevance representations, where each operation is called a ‘head.’ Eventually, a linear layer projects their concatenated outputs and summarizes the attention result.

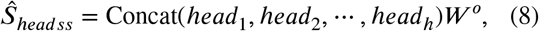

where *W °* is the head-wise weight matrix while a linear projection is applied after the output of the multi-head attention for each round.

As introduced above, the output of the multi-head layer is fed to the MLP layers, with GELU being their activation function. Simultaneously, a residual connection skips each MLP and connects to the output of the building block to avoid gradient vanishing. For downstream tasks, we solely use *S*_Cls_, which contains spatio-temporal-channel information for adaptive decision-making such as classification or other tasks. We can explicitly construct a correspondence between the input sEMG samples and different movements in the classification problem so the Transformer outputs probabilities of different classes.

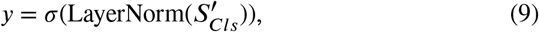

where *y* denotes the different targets of sEMG recognition, and 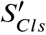 is the learned class token that is also normalized before the final decision-making module.

## 4. Experiments

### 4.1 Database

This section briefly discusses the two publicly available benchmark datasets used in the experiments, that is, NinaPro DB2 and NinaPro DB3 from offical Ninaweb^1^. In the respective measurement sessions, the sEMG sensors were placed on various muscle locations on the upper limbs. The datasets consist of hand activities broadly categorized into gestures, wrist movements, grasping objects, hand movements, etc.

#### NinaPro DB2

In DB2, the EMG data were collected from 40 healthy subjects (12 females, 6 left-handed and average aged 30 years) who performed in total 49 movements (8 isometric and isotonic hand configurations, 9 basic wrist movements, 23 grasping and functional movements and 9 force patterns). These movements are considered relevant to daily activities of living. These 49 movements are further divided into 4 groups, and each movement was repeated 6 times with a 3-second rest period between. The sEMG signal was recorded using 12 electrodes of a Delsys Trigno Wireless system, which provides a sampling rate of 2,000 Hz [24].

#### NinaPro DB3

The DB3 contains 11 experimental subjects who are the trans-radial amputees, other information is exactly the same as NinaPro-DB2. According to the authors of NinaPro database, three amputated subjects performed only a part of gestures due to fatigue or pain (3 subjects), and in 2 amputated subjects, the number of electrodes was reduced to ten due to insufficient space, hence the sEMG recordings from 6 six subject are used in this work following the experimental configuration used by [16].

### 4.2. Experimental Setup

Experiments are conducted by the PyTorch deep learning framework version 1.5.0. We used the `AdamW` optimizer to minimize the `cross entropy` loss function for model training. We perform a record-wise 5-fold cross-validation for the model: in each trial, 80% (32 subjects in DB2 and 5 subjects in DB3) of the recordings of samples are used to train models with the other (8 and 1 subjects, respectively) used for the validations. We summarize all parameters in Table 2.

**Table 2.**
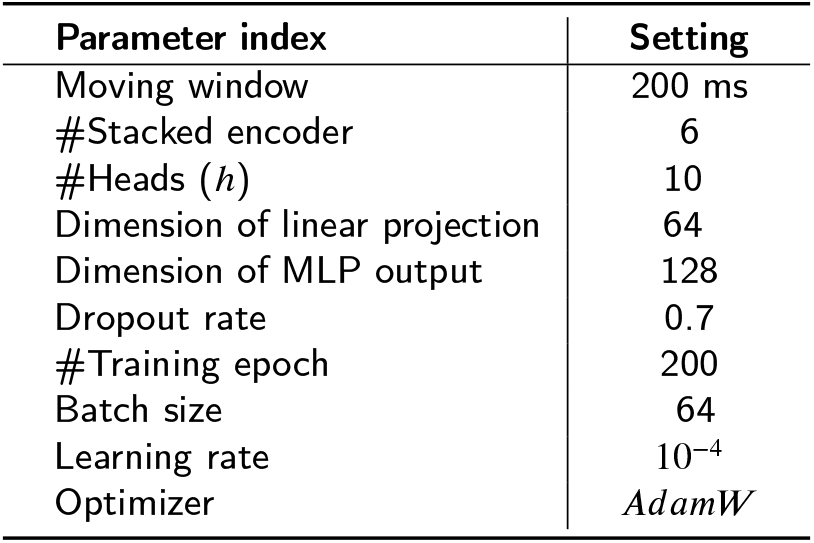
Parameter settings in this study.

| Parameter index | Setting |
| --- | --- |
| Moving window | 200 ms |
| #Stacked encoder | 6 |
| #Heads ($h$) | 10 |
| Dimension of linear projection | 64 |
| Dimension of MLP output | 128 |
| Dropout rate | 0.7 |
| #Training epoch | 200 |
| Batch size | 64 |
| Learning rate | $10^{-4}$ |
| Optimizer | *AdamW* |

### 4.3. Ablation Study

Extensive ablation studies are conducted to demonstrate how the proposed method performs when removing specific components. Since we advocate for channel-wise Transformer, we compare the proposed method against:

- *Temporal-CNN with/without Transformer*: CNNs with different kernels are utilized to summarize the temporal information from the raw sEMG signal.
- *Temporal-CNN + LSTM*: Temporal-CNN is associated with the LSTM to model the sequential order information.
- *Wavelet + LSTM*: LSTM is designed on top of DTCWT (referring to Section 3.1) to extract the sequential information in the time domain.
- *Feature-vector-wise Transformer*: The input of Transformer is the different feature index of spatio-temporal feature processed in Section 3.3, and each feature index summarizes all channel information.
- *Channel-wise LSTM/bi-LSTM*: LSTM/bi-LSTM replace to the Transformer that aims to evaluate the effectiveness of whether the parallel attention mechanism is better to the sequential order.

## 5. Results and Discussion

### 5.1. Results

#### *Observation 1:* The proposed model enjoys fast and smooth learning

From Figure 4 it is visible that the proposed method enjoys fast learning. The accuracy lines in Figure 4 (a) quickly reached above 0.8 and 0.6 respectively within only 25 epochs. Compared to the convergence rate in the most of previous studies (normally after 50-75), the proposed method has the power in summarizing the representational features for downstream tasks. Moreover, from the right subplot in Figure 4 (b) it can be seen that the loss curves are smooth, even though the training epoch (200) is far from the 25 iterations. The convergence range in DB2 and DB3 has a consistency after the training, such results indicating sufficient training of the model without overfitting issues.

**Figure 4:**
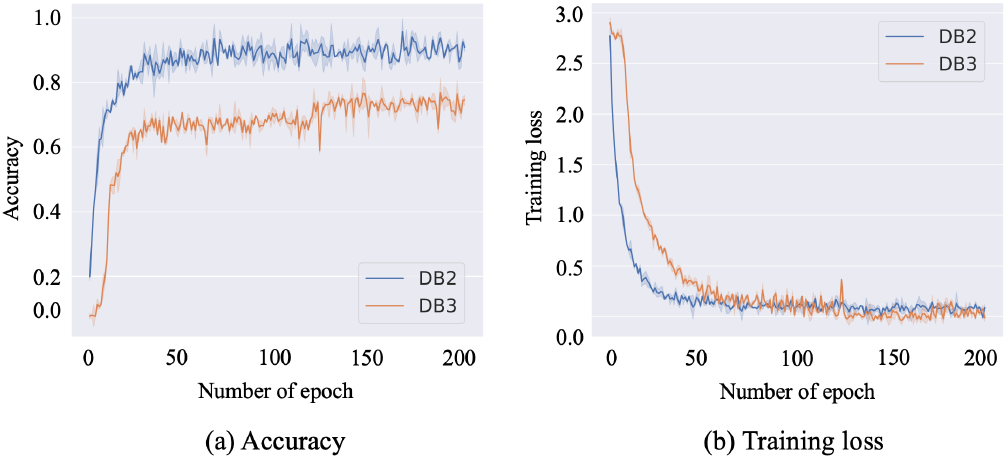
Accuracy and training loss of the proposed model on Ninapro DB2 and DB3.

#### Observation 2: The proposed model achieves SOTA and generalization ability

We compared the proposed method against existing state-of-the-art (SOTA) methods on both Ninapro DB2 and DB3 dataset and summarized the results in Table 3 and Table 4. It is visible that the proposed framework achieved new SOTA on both datasets. As we can see, the proposal achieves 0.91 accuracy which outperforms the previous work in DB2. Although the performance of the proposed method in sole DB3 does not reach the best result, applying the pretrain-to-fine-tuning strategy based on DB2, the proposed method has the best accuracy which outperforms the 0.81 accuracy in work [47]. However, both training and testing accuracy on DB3 dataset seems to underperform that of DB2. This is because DB3 consists of a small amount of samples recorded from amputees. To improve the performance, we pretrain the model on DB2 and then transfer it to DB3 for fine-tuning. The resulting performance reached an accuracy of 0.84, which proves the superiority of the model as well as its generalization ability across datasets.

**Table 3.**
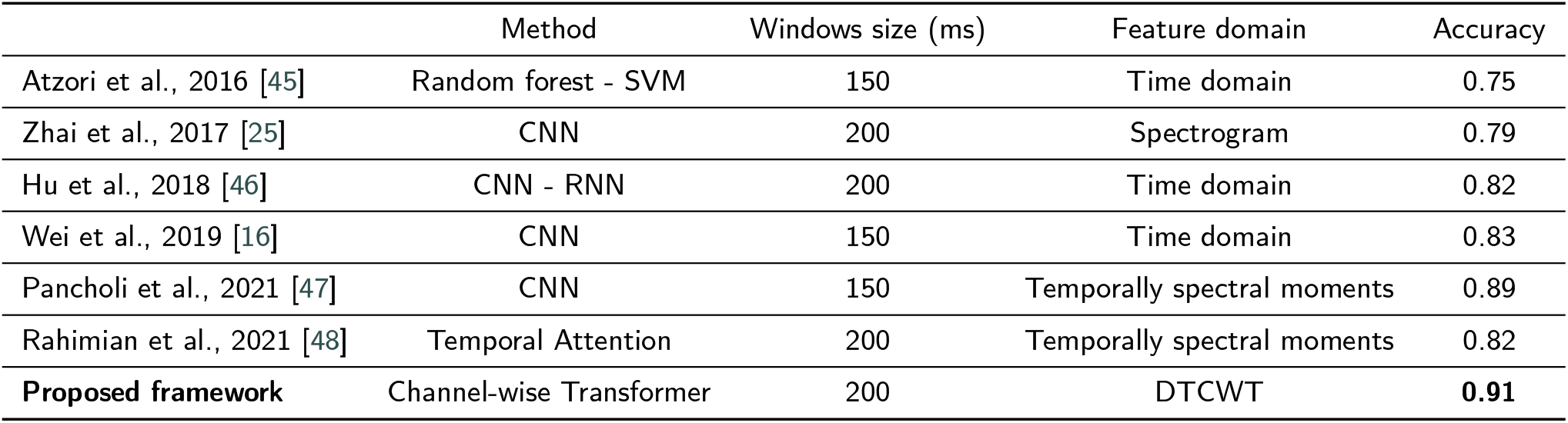
Comparison of the performance of the proposed model against existing state-of-the-art methods on the DB2 database.

**Table 4.** Performance obtained by the proposed model and existing methods using DB3 database.

|  | Method | Windows size (ms) | Feature domain | Accuracy |
| --- | --- | --- | --- | --- |
| Atzori et al., 2016 [45] | Random forest - SVM | 150 | Time domain | 0.46 |
| Zhai et al., 2017 [25] | CNN | 200 | Spectrogram | 0.73 |
| Wei et al., 2019 [16] | CNN | 150 | Time domain | 0.64 |
| Pancholi et al., 2021 [47] | CNN | 150 | Temporally spectral moments | 0.81 |
| Proposed framework | Channel-wise Transformer | 200 | DTCWT | 0.77 |
| **Proposed framework** | Channel-wise Transformer (Pre-trained DB3) | 200 | DTCWT | **0.84** |

#### *Observation 3:* Different window sizes and the number of features have an impact on the recognition performance of sEMG

In Table 3 and Table 4 we see the related work used different window sizes. To verify which size empirically works the best for the proposed model we swept the set [100, 150, 200, 400, 800] on both datasets. From Figure 5 it is visible that window size 200 performed the best on both DB2 and DB3. On the other hand, overly large window sizes have a negative impact on the performance since they might include more redundant noises.

**Figure 5:**
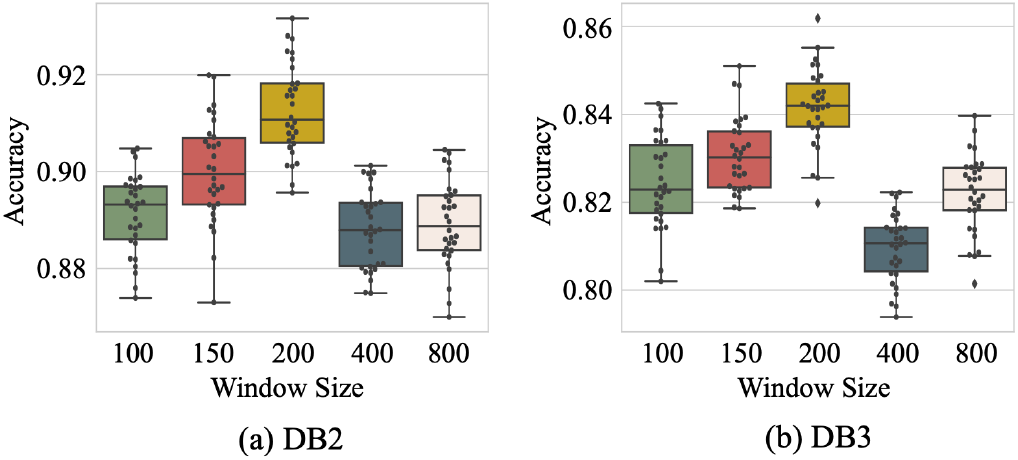
Ablation study of the impact of window size on accuracy. The window size within 200 has the best performance.

Since we introduce the MLP dropout feature ranking into this work, Figure 6 shows classification performances in the different feature vector settings with different numbers of features. With the decreasing number of features, the performance has gradually improved. Table 5 shows the details of selected features. Finally, only 8 features are the representative components embedded in sEMG channels used in this work. However, the classification performance is reduced when the *F4* (Kurtosis) and *F8* (Maximum Fractal Length) have dropped out of the feature vector, that is, these two features might have strong correlations to the sEMG recognition. This result proves there is some redundant and alternative information in the feature extraction or selection.

**Table 5.**
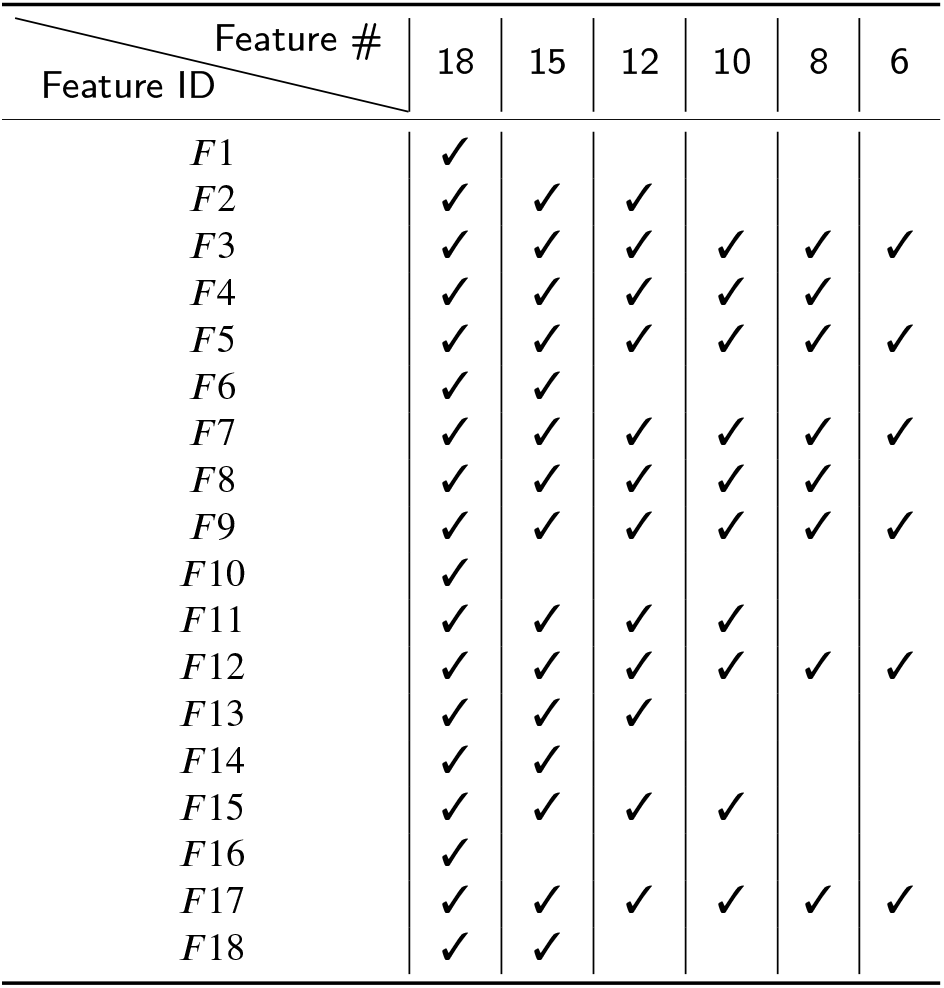
Selected features with each number of features.

**Figure 6:**
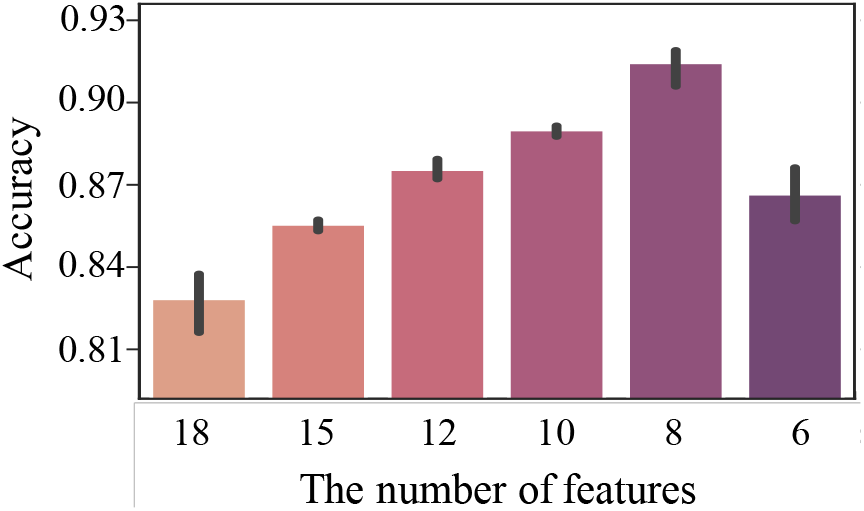
Feature reduction by MLP feature dropout method and the classification performance in DB2.

#### *Observation 4:* The Transformer outperforms other architectures on the investigated problems

In Figure 7 we compared the performance of various ablation choices introduced in Section 4.3. The five red columns show the accuracy resulting from different feature extraction methods. However, they are all significantly lower than the two green columns representing LSTM-based channel-wise methods. Nevertheless, the green columns are also outperformed by Transformer by around 0.1, indicating the limitation brought by the temporal and Markov properties of the LSTM. As summary, it might be safe to conclude that the Transformer architecture achieves the best among the discussed candidates on the investigated problems.

**Figure 7:**
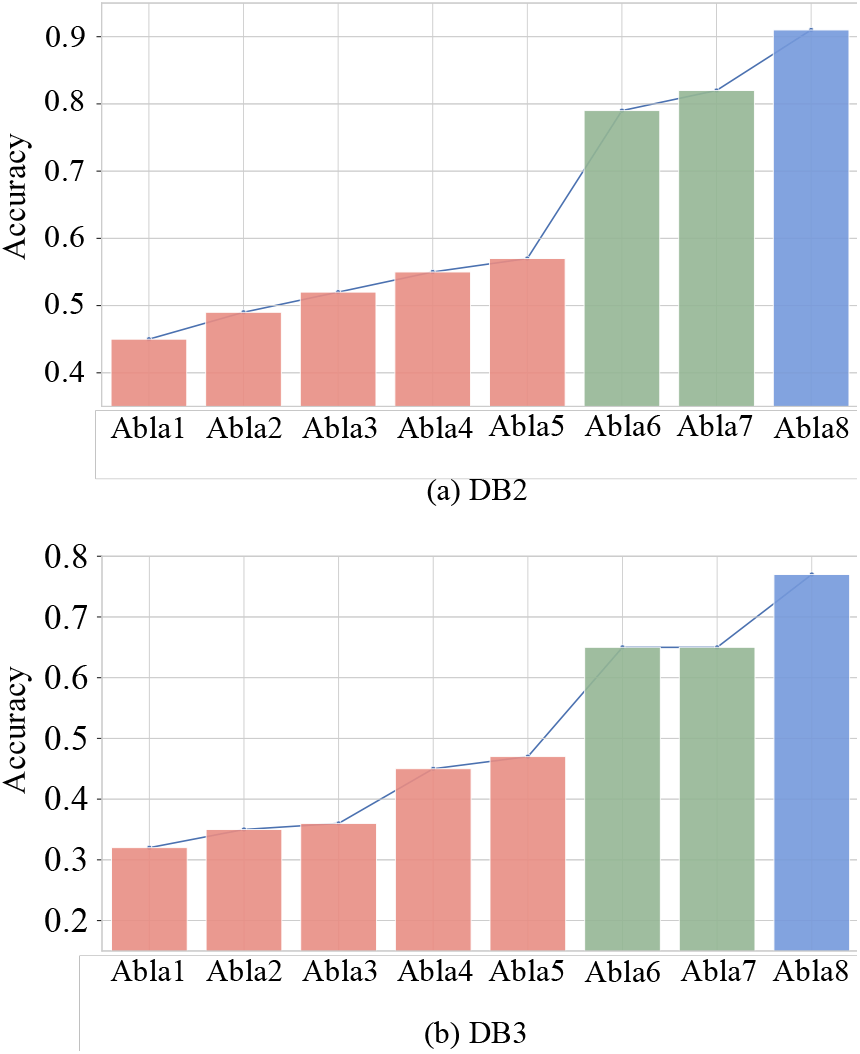
Performance of Ablation studies. Abla1: Temporal-CNN; Abla2: Temporal-CNN+LSTM; Abla3: Wavelet+LSTM; Abla4: Temporal-CNN+Transformer; Abla5: Feature-wise+Transformer; Abla6: Unidirectional LSTM; Abla7: Bi-directional LSTM; Abla8 (Proposal): Channel-wise Transformer

### 5.2. Discussion and future work

We have to point out that this study is based on preprocessing of spatio-temporal features: DTCWT and different statistical feature extraction methods and the real input of the Transformer model is a processed feature vector, hence the proposal is a semi-automatic sEMG recognition framework. The feature vector generation inevitably derives some costs in either time or computation. This is a limitation that extends our proposal to the tiny device or IoT environment since the real-time environment requires the lower computational costs possible. One potential solution is to consider an end-to-end deep learning architecture that uses different deep learning methods to replace such conventional feature extractions and to generate the same quality features by automatic findings. Future work will make efforts in this point to explore good candidates for model designs. However, a de novo deep network sometimes has a large model size that is also limited to the tiny device. The main reason is the large-scale parameters, hence, a good model quantization is also needed to transfer the model to a real-time IoT setting. Although the performance in gesture classification tasks based on public datasets is promising, the work lacks more general validation in different sEMG tasks. Considering a general-purpose model, we will find more suitable experiments for more sEMG tasks, such as robotics control or disorder environments. Meanwhile, more reliable evaluations in non-public/private datasets have been listed in our future plan.

## 6. Conclusion

In this paper, we proposed a novel 3-view sEMG feature representation enforced by conventional feature processing as well as the state-of-the-art Transformer. As proof of concept, sEMG classification was considered in this work: on two datasets consisting of both healthy people and amputees, we verified the proposed method attained state-of-the-art accuracy compared to existing methods. We also found the proposed model enjoys strong generalization ability in that the highest accuracy on the amputee dataset was achieved by pretraining the model on the healthy people data. We believe this time-frequency-channel representation is generally applicable and leveraging it to studies on gaits is one interesting future direction.

## Acknowledgment

This research and development work was partly supported by JSPS KAKENHI JP21K19828.

## Footnotes

1 http://ninaweb.hevs.ch/node/7

